# Temperature-dependent genetics of thermotolerance between yeast species

**DOI:** 10.1101/2022.01.11.475859

**Authors:** Melanie B. Abrams, Rachel B. Brem

## Abstract

Many traits of industrial and basic biological interest arose long ago, and manifest now as fixed differences between a focal species and its reproductively isolated relatives. In these systems, extant individuals can hold clues to the mechanisms by which phenotypes evolved in their ancestors. We harnessed yeast thermotolerance as a test case for such molecular-genetic inferences. In viability experiments, we showed that extant *Saccharomyces* cerevisiae survived at temperatures where cultures of its sister species *S. paradoxus* died out. Then, focusing on loci that contribute to this difference, we found that the genetic mechanisms of high-temperature growth changed with temperature. We also uncovered an enrichment of low-frequency variants at thermotolerance loci in *S. cerevisiae* population sequences, suggestive of a history of non-neutral selective forces acting at these genes. We interpret our results in light of a model of gradual acquisition of thermotolerance in the *S. cerevisiae* lineage by positive selection along a temperature cline. We propose that in an ancestral *S. cerevisiae* population, alleles conferring defects at a given temperature would have been resolved by adaptive mutations, expanding the range and setting the stage for further temperature advances. Together, our results and interpretation underscore the power of genetic approaches to explore how an ancient trait came to be.

## 1 Introduction

A central goal of evolutionary genetics is to understand how nature builds new phenotypes. Thanks to advances in statistical genetics and experimental evolution, mechanisms of trait evolution over relatively short timescales have come well within reach in the modern literature. By contrast, longer-term innovations have posed a more profound challenge for the field (Orr, 2001). In principle, for a phenotype that originated long ago and manifests now as a fixed difference between species, evolution could have refined the character along the entire divergence time of the respective taxa. In landmark cases, candidate-gene studies have shed light on suites of mutational changes of this kind between species, at a given model locus. This includes the order by which adaptive alleles were likely acquired, and/or the functional pressures that drove them (Bridgham et al., 2009; Baldwin et al., 2014; Dong et al., 2014; Sayou et al., 2014; Anderson et al., 2015; Daugherty et al., 2016; Sulak et al., 2016; Liu et al., 2018; Starr et al., 2018; Xie et al., 2018; Siddiq and Thornton, 2019; Pillai et al., 2020; Prieto-Godino et al., 2021). But, in most cases, any factor pursued by such a candidate-gene approach only represents part of the complex genetic architecture of an ancient trait. We still know relatively little about how evolution coordinates multiple adaptive loci over deep divergences.

In the search for evolutionary principles, genetically tractable model systems can be of great utility. *Saccharomyces* yeasts are well suited for this purpose, and environmental yeast isolates have been studied extensively for their innovations within and between species (Hittinger, 2013). Thermotolerance is of particular interest because it tracks with phylogeny across the 20 million years of the *Saccharomyces* radiation (Gonçalves et al., 2011; Salvadó et al., 2011). Even the two most recent branches of the phylogeny exhibit a robust difference in this phenotype: *S. cerevisiae* acquired the ability to grow at temperatures near 40°C in the five million years since it diverged from its sister species, *S. paradoxus* (Sweeney et al., 2004; Gonçalves et al., 2011; Salvadó et al., 2011; Williams et al., 2015). Previously, a whole-genome mapping scheme was used to identify housekeeping genes at which variation between *S. cerevisiae* and *S. paradoxus* impacts growth at the high end of the *S. cerevisiae* temperature range (Weiss et al., 2018). These loci exhibit striking sequence differences between the species, and conservation in *S. cerevisiae*, consistent with a history of positive selection on pro-thermotolerance alleles (Weiss et al., 2018; Abrams et al., 2021a, 2021b). However, we have as yet little insight into the ecological dynamics by which this model trait evolved along the *S. cerevisiae* lineage.

Cases of adaptation across temperature clines are a mainstay of the evolutionary genetics literature (Turner et al., 2008; Prasad et al., 2011; Mimura et al., 2013; Savolainen et al., 2013; Robin et al., 2017; Dudaniec et al., 2018; Key et al., 2018; Endler, 2020; Tepolt and Palumbi, 2020; Calfee et al., 2021; Machado et al., 2021). Here we sought to explore the relevance of such a mechanism to the events by which *S. cerevisiae* gained its maximal thermotolerance phenotype. Given that we have no access to genotypes representing ancient intermediates between this species and *S. paradoxus*, we designed a strategy to interrogate the genetics of extant strains, focusing on contributing genes of major effect. We surveyed gene-environment interactions by these thermotolerance loci across warm temperatures, complementing previous studies of interspecies variation at a single high temperature (Weiss et al., 2018; Abrams et al., 2021b). And we investigated the frequency of variants at these loci with a population-genomic approach.

## 2 Methods

### 2.1 Dose response growth assay

For growth measurements in Figure 2, we assayed *S. paradoxus* Z1, S. cerevisiae DBVPG1373, and *S. cerevisiae* DBVPG1373 with the *S. paradoxus* Z1 allele of *ESP1, MYO1, AFG2*, or *CEP3* swapped in at the endogenous locus from (Weiss et al., 2018) (Table S2) as follows. Each strain was streaked from a -80°C freezer stock onto a yeast peptone dextrose (YPD) agar plate and incubated at room temperature for 3 days. For each biological replicate, a single colony was inoculated into 5 mL liquid YPD and grown for 24 hours at 28°C with shaking at 200 rpm to generate pre-cultures. Each pre-culture was backdiluted into YPD at an OD_600_/mL of 0.05 and grown for an additional 5.5-6 hours at 28°C, shaking at 200 rpm, until reaching logarithmic phase. Each pre-culture was again back-diluted into 10 mL YPD in 1-inch diameter glass tubes with a target OD_600_/mL of 0.05; the actual OD_600_/mL of each was measured, after which it was grown at a temperature of interest (28°C or 35-38°C) with shaking at 200rpm for 24 hours, and OD_600_/mL was measured again. The growth efficiency for each replicate was calculated as the difference between these final and initial OD_600_/mL values. We used the growth efficiency from all days and all temperatures of a given strain s as input into a two-factor type 2 ANOVA test for a temperature-by-strain effect comparing s with *S. cerevisiae*. For growth measurements of *ESP1* swap strains with different donors, as reported in Figure S2, cultures were grown and measured as above, except that the only temperature was 36°C.

### 2.2 Viability assay

For the survey of viability phenotypes at high temperatures across wild-type isolates in Figure 2, strains were streaked out and a colony of each was pre-cultured in liquid as for 39°C growth above, except that the initial pre-culture to achieve saturation lasted 48 hours. Each pre-culture was back-diluted into 10 mL of YPD to reach an OD_600_/mL of 0.05 and then cultured for 24 hours at the temperature of interest (28°C - 39°C). The viability of both the precultures and the cultures after 24 hours at the temperature of interest were measured with a spotting assay, where we diluted aliquots from the culture in a 1:10 series and spotting 3 µL of each dilution for growth on a solid YPD plate. After incubation at 28°C for two days, we used the dilution corresponding to the densest spot that was not a lawn for to determine viability: we counted the number of colonies in each of the two technical replicate spots, formulated the number of colony forming units per mL of undiluted culture (CFU/mL). We determined the change in viable cells by subtracting the number of cells in the culture at the initial time point from that at the final timepoint, based on the CFU/mL count and the culture volume. We evaluated the significance of the difference between *S. cerevisiae* and *S. paradoxus* at a given temperature using a one-sided Mann-Whitney U test.

### 2.3 Tajima’s D in Wine/European *S. cerevisiae*

Tajima’s D tabulates the difference between the average number of differences in pairs of sequences in a population sample and the number of variant sites in the sample. When the latter is of much bigger magnitude and D is negative, it indicates that variation in the sample is accounted for mostly by rare alleles. This pattern is expected some time after a selective sweep when de novo mutations arise on the swept haplotype; it can also reflect weak purifying selection or a history of population expansion (Suzuki, 2010). Taking as input the VCFs reporting inheritance in the strains of a given *S. cerevisiae* population from (Peter et al., 2018), we used VCF-kit (Cook and Andersen, 2017) to calculate Tajima’s D for the region from coding start to coding stop for each gene. We then developed a resampling test to assess enrichment trends in Tajima’s D in genes of interest against a genomic null. This scheme normalizes out impacts on Tajima’s D values from events which affect the whole genome, such as a population expansion; that said, formally, any results from such an empirical outlier-based analysis serve as suggestive rather than conclusive evidence for non-neutral evolution (Teshima et al., 2006; Thornton and Jensen, 2007). For the test, we sampled 10,000 random cohorts of genes from the genome with the same number of essential and nonessential genes as our thermotolerance cohort (Winzeler et al., 1999), and we used as an empirical P-value the proportion of random cohorts where the median Tajima’s D was less than or equal to that of our thermotolerance cohort. Application of this test to the Wine/European *S. cerevisiae* population (362 strains) is reported in Figure 3 of the main text. Applied to the Mosaic Region 3 (113 strains), Mixed Origin (72 strains), Sake (47 strains), and Brazilian Bioethanol (35 strains), the four next most deeply sampled populations from (Peter et al., 2018), this test for enrichment of low Tajima’s D among our four focal thermotolerance genes yielded *P* = 0.8351, 0.975, 0.8624, and 0.0348 respectively.

## 3 Results

For an initial study of the genetics of yeast species variation in temperature response, we chose to harness DBVPG1373, an *S. cerevisiae* isolate from Dutch soil, and Z1, an *S. paradoxus* isolate from an oak tree in England. We anticipated that detailed analyses using these strains, as representatives of their respective species, could accelerate the discovery of more general principles (Weiss et al., 2018; Abrams et al., 2021a, 2021b). We developed an assay quantifying cell viability in a given liquid culture before and after incubation at a temperature of interest, and we implemented this approach for each species in turn. The results revealed an advantage for *S. cerevisiae* over *S. paradoxus* at temperatures above 35°C (Figure 1), consistent with previous growth-based surveys (Sweeney et al., 2004; Gonçalves et al., 2011; Salvadó et al., 2011; Williams et al., 2015). *S. cerevisiae* maintained viability at all temperatures tested, whereas by 37°C, *S. paradoxus* became inviable, with no evidence for spontaneous rescue even over long incubation times (Figure 1). Using the latter as a window onto the phenotype of the common ancestor of *S. cerevisiae* and *S. paradoxus*, we would envision that the latter ancient population would have gone extinct rather than adapting, if exposed to temperatures at the high end of the range tolerated by modern *S. cerevisiae*.

**Figure 1.**
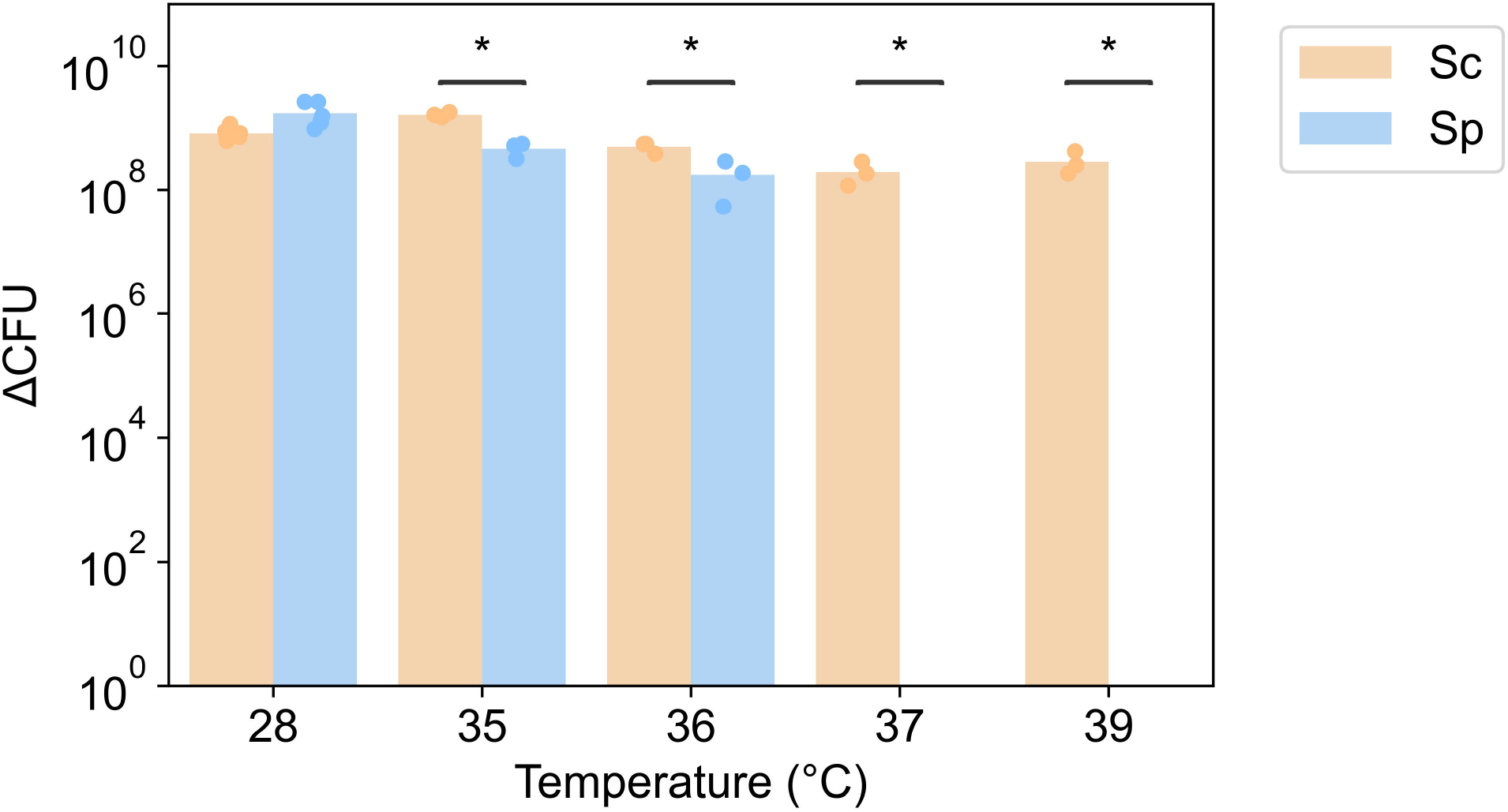
*S. paradoxus* is inviable at high temperature. Each column reports the viability of a culture of *S. cerevisiae* DBVPG1373 or *S. paradoxus* Z1 at the indicated temperature. The *y*-axis reports the number of viable colony forming units (CFU) per unit of optical density after 24 hours of incubation, as a difference from the analogous quantity at time zero. Each point reports results from one biological replicate, and each bar height reports the average across replicates (*n* = 3-6). *, *P* < 0.05 for a one-sided Mann-Whitney U test assessing the advantage of *S. cerevisiae* over *S. paradoxus* at a given temperature.

We next aimed to investigate the genetics of temperature response as it differs between extant *S. cerevisiae* and *S. paradoxus*, to motivate inferences about the evolution of the maximal thermotolerance trait. We focused on four genes—the cell division factors *ESP1, MYO1*, and *CEP3*, and the ribosome maturation factor *AFG2*—where alleles from modern *S. paradoxus* compromise growth at 39°C (Weiss et al., 2018). We made use of strains of the *S. cerevisiae* DBVPG1373 background harboring the allele of each gene in turn from *S. paradoxus* Z1. In each, we measured the growth phenotype as a dose-response across temperatures, and observed a drop as temperature increased (Figure 2). In the *S. cerevisiae* background, *S. paradoxus* Z1 alleles eroded growth efficiency at temperatures well below the hard limit of viability for the Z1 wild-type (∼38°C), an effect that reached significance for three of our four focal genes (Figure 2). We conclude that many problems posed by *S. paradoxus* Z1 alleles at the high end of our temperature range also manifest to a lesser extent at lower temperatures.

**Figure 2.**
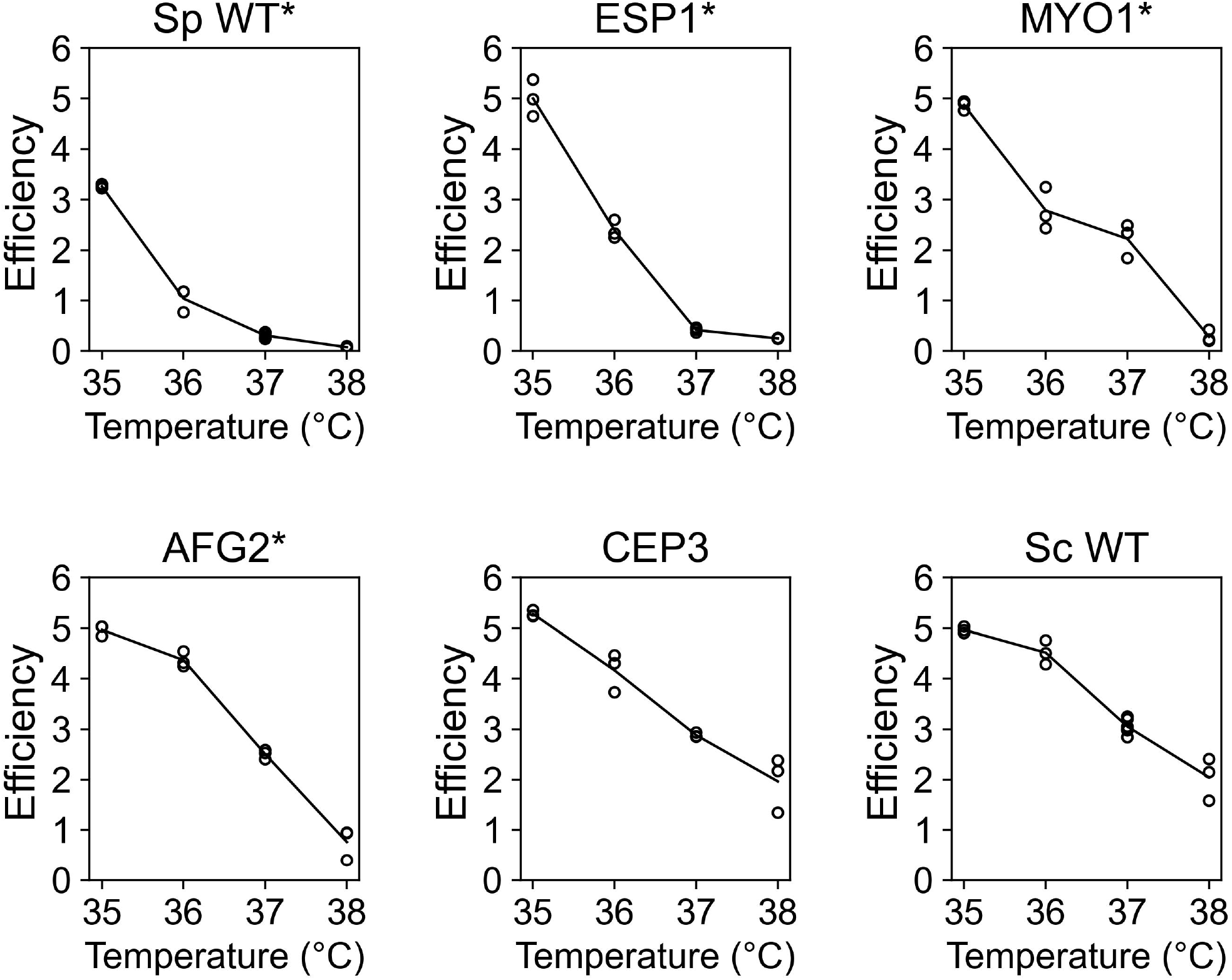
*S. paradoxus* alleles of thermotolerance genes confer temperature-dependent defects in the *S. cerevisiae* background. Each panel reports the results of growth experiments of a strain of *S. cerevisiae* DBVPG1373 harboring the indicated gene from *S. paradoxus* Z1, or the respective wild-type parent strains (WT), across a temperature gradient. In a given panel, the *y*-axis reports growth efficiency, the optical density reached by the culture after 24 hours at the temperature indicated on the *x*-axis, as a difference from the analogous quantity at time zero. Each point reports results from one biological replicate (*n* = 3), and the line represents the average growth efficiency of the indicated strain across the temperature gradient. *, *P*□<□10^−3^ for the strain by temperature interaction term of a two-factor ANOVA, in a comparison between the indicated strain and wild-type *S. cerevisiae*.

Inspecting the shape of the temperature dose-responses of allelic effects, we noted quantitative differences between our loci. At the chromatid separase gene *ESP1*, the Z1 allele conferred an appreciable drop in growth efficiency at 36°C and supported almost no growth by 37°C (Figure 2B). By contrast, at *AFG2*, the *S. paradoxus* Z1 allele was sufficient for growth approximating that of *S. cerevisiae* until 38°C (Figure 2D). The dose-response of allelic effects at *MYO1*, encoding a class II myosin heavy chain, lay between these two extremes (Figure 2C). This differential susceptibility to temperature across the genes of our set likely reflects distinguishing properties of their structure and function, and of the interspecies variants they harbor.

We reasoned that trends from our temperature dose-response approach would be most informative when they were conserved across *S. paradoxus* as a species. To pursue this, we earmarked *ESP1*, whose *S. paradoxus* Z1 allele had exhibited the sharpest falloff with temperature among the genes of our set (Figure 2B). We repeated our growth efficiency experiments in strains of *S. cerevisiae* DBVPG1373 harboring *ESP1* from other *S. paradoxus* donors beside the Z1 strain. These transgenics, which phenocopy wild-type DBVPG1373 at 28°C (Weiss et al., 2018), all dropped off in growth efficiency by 36°C (Figure S2), as expected if the temperature preference of *ESP1* were ancestral to, and shared across, extant *S. paradoxus* populations.

Together, these dose-response results make clear that the functions of *S. paradoxus* alleles at our focal genes break down at distinct temperatures between 35°C and 38°C—suggesting similar gene-by-environment effects in the ancestor of *S. cerevisiae* and *S. paradoxus*, if it sampled a range of temperature conditions over evolutionary history.

To gain insight into past and current selective pressures on thermotolerance loci, we turned to a molecular-evolution approach. Previous work on *Saccharomyces* thermotolerance genes has emphasized sequence divergence between species (Weiss et al., 2018; Abrams et al., 2021a, 2021b). For a complementary focus on population variation within *S. cerevisiae*, we analyzed the Tajima’s D statistic, which formulates properties of sequence variants in a population into a test of how well the sequence fits the expectations under a neutral evolutionary model (Tajima, 1989; Biswas and Akey, 2006). We first developed a resampling-based scheme that compares Tajima’s D between a gene cohort of interest and a genomic null. This enables a non-parametric assessment of significance and normalizes out potential effects of demography on the statistic (see Methods). Expecting that our test would have maximal power in a data set of large sample size, we focused on the most deeply-sampled *S. cerevisiae* population in current compendia, comprising isolates from vineyards and European soil (Peter et al., 2018). Examining our four focal thermotolerance genes, we detected an enrichment for low, negative Tajima’s D at these loci across *S. cerevisiae* genomes (Figure 3 and Table S1), reporting an excess of rare variants—as expected after a selective sweep, or under constraints from purifying selection (Biswas and Akey, 2006; Suzuki, 2010). We repeated this analysis using more comprehensive sets of hits from interspecies thermotolerance screens (Weiss et al., 2018; Abrams et al., 2021a, 2021b), and detected strong signal for low, negative Tajima’s D at these loci in vineyard/European *S. cerevisiae* in every case (Table S1). Interestingly, *ESP1* exhibited the most negative Tajima’s D value among all thermotolerance genes (Table S1), dovetailing with the strong effect of variation at this gene in phenotypic analyses (Weiss et al. 2018 and Figure 2). The trend for low, negative Tajima’s D in thermotolerance genes was detectable but not consistent across other shallowly-sampled populations of *S. cerevisiae* (see Methods), potentially reflecting weaker power or weaker selection in the latter relative to vineyard/European strains. In either case, our data establish that thermotolerance gene variation in some modern *S. cerevisiae* populations is consistent with a history of non-neutral evolution.

**Figure 3.**
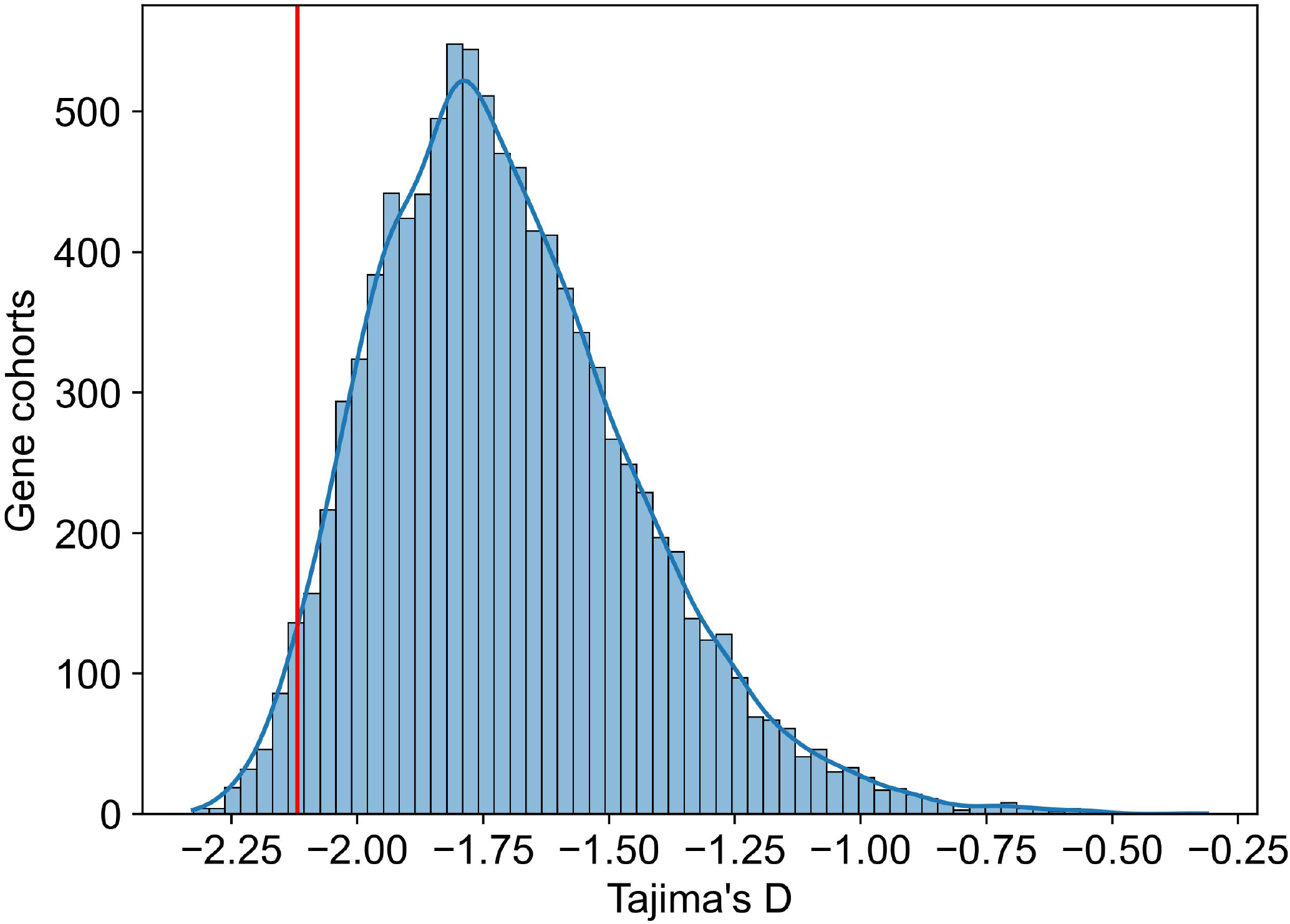
Thermotolerance genes are enriched for low Tajima’s D in *S. cerevisiae*. The *x*-axis reports the median Tajima’s D across a gene set of interest in genomes of wine/European strains of *S. cerevisiae* from (Peter et al., 2018). Blue bars reflect the results from genomic resampling, with the *y*-axis reporting the number of randomly chosen gene sets with the median Tajima’s D shown on the *x*, and the blue curve showing a kernel density estimate of the histogram bar values. The red vertical line reports the Tajima’s D value of the four thermotolerance genes characterized in Figure 2, corresponding to resampling *P* = 0.0263.

## 4 Discussion

Most trait differences that define long-diverged species are likely the product of suites of unlinked variants that have come together over long timescales. Tracing the ancient evolutionary events at such loci remains a central challenge in the field. In this work, we have characterized thermotolerance genes in extant yeast strains and species to inform models of the evolution of the trait in *S. cerevisiae*.

Our data have shown that temperatures below the high end of the *S. cerevisiae* range are lethal for *S. paradoxus*. This complements previous surveys of the *Saccharomyces* clade using growth-based assays (Sweeney et al., 2004; Gonçalves et al., 2011; Salvadó et al., 2011) and reveals that, under high temperature conditions, a given culture of *S. paradoxus* will die off rather than adapting. Assuming similar behavior in an ancestor of *S. cerevisiae* before the rise of its modern thermotolerance profile, we infer that the latter event was unlikely to be precipitated by a single jump to a hot growth environment, long ago in history. Rather, we favor the hypothesis that thermotolerance evolution through the *S. cerevisiae* lineage proceeded along a temperature cline, as has been documented in elegant local-adaptation case studies (Mimura et al., 2013; Robin et al., 2017; Dudaniec et al., 2018; Key et al., 2018; Tepolt and Palumbi, 2020). In this scenario, a well-adapted ancestral *S. cerevisiae* population in a given temperature niche would have acquired variants that resolved defects manifesting in slightly warmer conditions, and expanded its range. If the *S. cerevisiae* lineage did go through a series of such transitions in acquiring maximal thermotolerance, it would echo the principle that gradual exposure to increasing stresses fosters adaptation and reduces the risk of extinction, relative to a sudden, high dose of stress (Lindsey et al., 2013).

We have also leveraged results from genetic mapping of *S. cerevisiae* thermotolerance to trace the temperature dependence of contributing genes. This contrasts with approaches that search genomes for variants associated with environment or population variables (Rellstab et al., 2015; Hoban et al., 2016), in that we focus directly on bona fide causal determinants of the trait of interest. Our strategy has allowed us to discern differences between thermotolerance loci in terms of the temperatures at which alleles from modern *S. paradoxus* fail in their growth functions. In other words, the genetic mechanisms of growth change with temperature in this system. If such effects were at play in the *S. cerevisiae* ancestor, as it sampled conditions and niches during the acquisition of thermotolerance, each locus would have come under selective pressure at the temperature where its defect manifested. This phenomenon, by which the weakest point in the genetic network targeted by evolution changes as conditions change, has been termed “ whack-a-mole” dynamics (Shin and MacCarthy, 2015). Ultimately, a complete model of adaptation across time and environment will take into account the shifts in the evolutionary landscape from this effect as well as from constant changes in genetic background (Starr and Thornton, 2016).

If the ancient *S. cerevisiae* population did adapt progressively along a temperature cline, what would the ecology have been? In principle, migrants from temperate physical locales could have advanced to warmer and warmer locales, perhaps terminating in the hot East Asian environments to which the ancestor of modern *S. cerevisiae* has been traced (Peter et al., 2018). Alternatively, the trait syndrome in this species—a unique ability to tolerate ethanol as well as heat, with both given off by fermentative metabolism—could have arisen as a specialization to kill off microbial competitors in nutrient-rich substrates, regardless of the endemic temperature of the location (Goddard, 2008; Salvadó et al., 2011). If so, variants would have been acquired, potentially over millions of generations, gradually to ratchet up fermentative activity and tolerance to its byproducts, with our genetic insights thus far largely restricted to the latter.

Our work also leaves open the dating of any such events. In several *S. cerevisiae* populations, we have uncovered an enrichment of rare variants at thermotolerance genes. This signature of non-neutral evolution can be interpreted at a given locus as evidence for a relatively recent sweep of a positively selected haplotype, or for negative selection to maintain a fitness-relevant haplotype that arose long ago. We favor the latter hypothesis, given that prior work across populations has also made clear that *S. cerevisiae* alleles at our focal genes are partly sufficient for maximal thermotolerance, conserved within the species, and divergent from *S. paradoxus* (Weiss et al., 2018; Abrams et al., 2021b). We thus propose that many thermotolerance alleles were acquired in ancient selective sweeps, before the radiation of modern *S. cerevisiae*, and have been maintained since then by purifying selection. That said, later refinements in particular populations of *S. cerevisiae* may also have strengthened beneficial facets of the trait, added regulatory tuning, or eliminated antagonistic pleiotropic “ side effects” that were niche-specific. A comprehensive genetic and ecological reconstruction of this history may be out of our current grasp, especially in light of the caveats of our approach using a laboratory setting and extant strain backgrounds. Nonetheless, our data add compelling detail to an emerging consensus view of how evolution built maximal thermotolerance in *S. cerevisiae*.

## 6 Author Contributions

M.A. and R.B.B. designed the study. MBA conducted experimental work and performed data collection and analysis. MBA and RBB wrote the manuscript.

## 8 Acknowledgments

The authors thank Faisal AlZaben, Abel Duarte, and Jude Edwards for experimental support and Adam Arkin for his generosity with computational resources.

## 9 Data Availability Statement

Strains and plasmids are available upon request. Custom scripts for the analysis will be available at https://github.com/melanieabrams-pub/.

