## Supplemental materials for "Temperature-dependent genetics of thermotolerance between yeast species"

Supplementary Material

### 1 Supplementary Figures and Tables

#### 1.1 Supplementary Figures

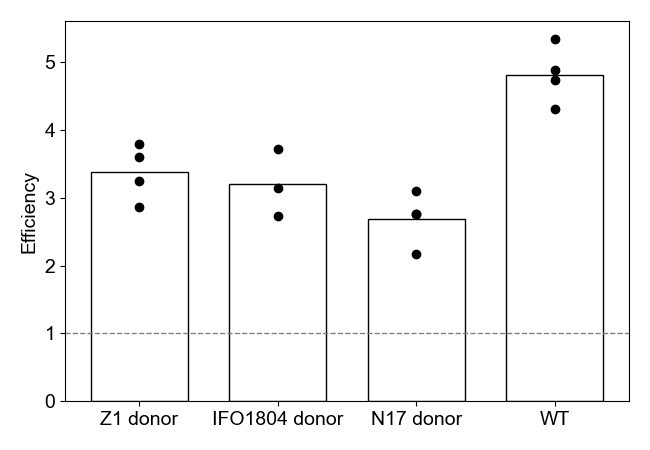

**Supplementary Figure 1. Alleles of *ESP1* from different *S. paradoxus* donor strains compromise high-temperature growth in the *S. cerevisiae* background.** Each column reports the results of growth experiments of a strain of *S. cerevisiae* DBVPG1373 harboring the *ESP1* allele from the indicated donor strain of *S. paradoxus*, or wild-type *S. cerevisiae* DBVPG1373 (WT). The *y*-axis reports the normalized growth efficiency at 36°C; in a given column, points report biological replicates (*n* = 3-4), and the bar height reports the average.

#### 1.2 Supplementary Tables

**(A)**

| **Gene** | **Common name** | **Tajima's D** |
| --- | --- | --- |
| *YGR098C* | *ESP1* | -2.390412929 |
| *YLR397C* | *AFG2* | -2.16261685 |
| *YHR023W* | *MYO1* | -2.075765368 |
| *YMR168C* | *CEP3* | -1.993925578 |
| Enrichment *P =* 0.0263 | | |

**(B)**

| **Gene** | **Common name** | **Tajima's D** |
| --- | --- | --- |
| *YGR098C* | *ESP1* | -2.390412929 |
| *YKR054C* | *DYN1* | -2.203644652 |
| *YLR397C* | *AFG2* | -2.16261685 |
| *YDR180W* | *SCC2* | -2.15471199 |
| *YHR023W* | *MYO1* | -2.075765368 |
| *YMR168C* | *CEP3* | -1.993925578 |
| *YNL172W* | *APC1* | -1.841207567 |
| *YCR042C* | *TAF2* | -1.791614606 |
| Enrichment *P* = 0.0046 | | |

**(C)**

| **Gene** | **Common name** | **Tajima's D** |
| --- | --- | --- |
| *YGR098C* | *ESP1* | -2.390412929 |
| *YKL134C* | *OCT1* | -2.293734473 |
| *YMR016C* | *SOK2* | -2.281861163 |
| *YKR054C* | *DYN1* | -2.203644652 |
| *YLR397C* | *AFG2* | -2.16261685 |
| *YDR180W* | *SCC2* | -2.15471199 |
| *YHR023W* | *MYO1* | -2.075765368 |
| *YDR443C* | *SSN2* | -2.070565734 |
| *YMR168C* | *CEP3* | -1.993925578 |
| *YPR164W* | *MMS1* | -1.990253695 |
| *YPL174C* | *NIP100* | -1.967856781 |
| *YCR042C* | *TAF2* | -1.791614606 |
| *YJR135C* | *MCM22* | -1.651398253 |
| *YJL025W* | *RRN7* | -1.502614461 |
| Enrichment *P* = 0.0010 | | |

**(D)**

| **Gene** | **Common name** | **Tajima's D** |
| --- | --- | --- |
| *YBR136W* | *MEC1* | -2.425141812 |
| *YMR275C* | *BUL1* | -2.421881839 |
| *YDR103W* | *STE5* | -2.336064982 |
| *YKL017C* | *HCS1* | -2.301392848 |
| *YGR140W* | *CBF2* | -2.268956348 |
| *YJR127C* | *RSF2* | -2.193879044 |
| *YNL132W* | *KRE33* | -2.185460334 |
| *YPL268W* | *PLC1* | -2.179277149 |
| *YLR397C* | *AFG2* | -2.16261685 |
| *YDR180W* | *SCC2* | -2.15471199 |
| *YKL197C* | *PEX1* | -2.146336148 |
| *YAL026C* | *DRS2* | -2.131356095 |
| *YGL082W* | *MIY1* | -2.118951228 |
| *YER151C* | *UBP3* | -2.07655222 |
| *YDL035C* | *GPR1* | -2.058601518 |
| *YDR508C* | *GNP1* | -2.023301799 |
| *YDR235W* | *PRP42* | -2.016907874 |
| *YLR422W* | *DCK1* | -2.009696117 |
| *YJL062W* | *LAS21* | -2.009120073 |
| *YIL068C* | *SEC6* | -1.946442537 |
| *YDR456W* | *NHX1* | -1.937222516 |
| *YGL095C* | *VPS45* | -1.936591083 |
| *YNR045W* | *PET494* | -1.908223035 |
| *YMR125W* | *STO1* | -1.833598797 |
| *YCR042C* | *TAF2* | -1.791614606 |
| *YNL049C* | *SFB2* | -1.786844048 |
| *YOR092W* | *ECM3* | -1.781980316 |
| *YML099C* | *ARG81* | -1.772396018 |
| *YOR326W* | *MYO2* | -1.746603341 |
| *YPL254W* | *HFI1* | -1.722539604 |
| *YJR107W* | *LIH1* | -1.718242115 |
| *YKL114C* | *APN1* | -1.709523846 |
| *YMR078C* | *CTF18* | -1.699771771 |
| *YPR049C* | *ATG11* | -1.696994126 |
| *YMR167W* | *MLH1* | -1.655296333 |
| *YOL081W* | *IRA2* | -1.568684518 |
| *YOR091W* | *TMA46* | -1.543998807 |
| *YMR207C* | *HFA1* | -1.220103519 |
| *YIL152W* | *VPR1* | -1.169622493 |
| *YMR094W* | *CTF13* | -1.159736527 |
| *YDR375C* | *BCS1* | -1.077792241 |
| *YDR318W* | *MCM21* | -0.780309448 |
| *YOR371C* | *GPB1* | -0.543345312 |
| *YGR198W* | *YPP1* | -0.451850166 |
| Enrichment *P =* 0.0012 | | |

**Supplementary Table 1. Genes from thermotolerance mapping are enriched for low Tajima’s D in *S. cerevisiae*.** Each panel reports the analysis, in one set of hit genes from genetic dissection of thermotolerance between *S. cerevisiae* and *S. paradoxus,* of Tajima’s D in genomes of wine/European strains of *S. cerevisiae* from (Peter et al. 2018). For a given panel, each of the first through penultimate rows reports the gene, common name, and Tajima’s D. The final row reports the significance of the enrichment of low Tajima’s D of the indicated genes in a genomic resampling test. (A) The four focal thermotolerance loci studied in this work. (B) The eight thermotolerance genes identified in (Weiss et al. 2018). (C) The 14 thermotolerance genes identified in (Abrams et al. 2021b). (D) The 44 thermotolerance genes identified in (Abrams et al. 2021a).

| **Species background** | **Strain background** | **Swap  (if applicable)** | **Source** | **Strain Name** |
| --- | --- | --- | --- | --- |
| *S. paradoxus* | Z1 | N/A | Weiss et al., 2018 | CW62 |
| *S. cerevisiae* | DBVPG1373 | N/A | Weiss et al., 2018 | CW68 |
| *S. cerevisiae* | DBVPG1373 | *S. paradoxus* Z1 CEP3 full swap | Weiss et al., 2018 | CW73 |
| *S. cerevisiae* | DBVPG1373 | *S. paradoxus* Z1 AFG2 full swap | Weiss et al., 2018 | CW64 |
| *S. cerevisiae* | DBVPG1373 | *S. paradoxus* Z1 MYO1 full swap | Weiss et al., 2018 | CW104 |
| *S. cerevisiae* | DBVPG1373 | *S. paradoxus* Z1 ESP1 full swap | Weiss et al., 2018 | CW98 |
| *S. cerevisiae* | DBVPG1373 | *S. paradoxus* N17 ESP1 full swap | Weiss et al., 2018 | CW284 |
| *S. cerevisiae* | DBVPG1373 | *S. paradoxus* IFO1804  ESP1 full swap | Weiss et al., 2018 | CW287 |

**Supplementary Table 2. Strains used in this study.**
